# Intersubject brain network organization during dynamic anxious anticipation

**DOI:** 10.1101/120451

**Authors:** Mahshid Najafi, Joshua Kinnison, Luiz Pessoa

## Abstract

How do large-scale brain networks reorganize during the waxing and waning of anxious anticipation? Here, threat was dynamically modulated during functional MRI as two circles slowly meandered on the screen; if they touched, an unpleasant shock was delivered. We employed intersubject network analysis, which allows the investigation of network-level properties “across brains,” and sought to determine how network properties changed during periods of approach (circles moving closer) and periods of retreat (circles moving apart). Dynamic threat altered network cohesion across the salience, executive, and task-negative networks, as well as subcortical regions. Functional connections between subcortical regions and the salience network also increased during approach vs. retreat, including the putative periaqueductal gray, habenula, and amygdala, showing that the latter is involved under conditions of relatively prolonged and uncertain threat (the bed nucleus of the stria terminalis was observed during both approach and retreat). Together, our findings unraveled dynamic properties of large-scale networks across participants while threat levels varied continuously, and demonstrate the potential of characterizing emotional processing at the level of distributed networks.

**Significance Statement:** Understanding the brain basis of anxious anticipation is important not only from a basic research perspective, but because aberrant responding to uncertain future negative events is believed to be central to anxiety disorders. Although previous studies have investigated how brain responses are sensitive to threat proximity, little is known about how patterns of response co-activation change during dynamic manipulations of threat. To address these important gaps in the literature, we studied the dynamics of emotional processing at the level of large-scale brain networks by devising a manipulation in which threat was dynamically modulated during functional MRI scanning.

## Introduction

Imagine yourself reclining on a dentist’s chair. Most of us experience an aversive reaction to the onset of the drill, and wait anxiously for the moment the dentist will finish checking it and move it toward our mouth. Understanding the brain basis of anxious anticipation is important not only from a basic research perspective, but because aberrant responding to uncertain future negative events is believed to be central to anxiety disorders (Grupe and Nitschke, 2013, Fox and Kalin, 2014). A growing literature of both non-human and human research indicates that anticipatory processing of negative events engages multiple brain regions (Davis et al., 2010a, Grupe and Nitschke, 2013, Tovote et al., 2015), including medial prefrontal cortex, insula, and orbitofrontal cortex, cortically. Subcortically, implicated regions include the amygdala, periaqueductal gray (PAG), and the bed nucleus of the stria terminalis (BNST); the latter has received considerable attention in the past decade (see Davis et al., 2010b, Fox et al., 2015).

Despite recent progress, important questions remain largely unanswered. First, how does anxious anticipation engage and reorganize large-scale brain networks? We propose that emotional processing needs to be characterized at the level of distributed networks (Pessoa, in press), and not just at the level of evoked responses in specific brain regions (such as the amygdala or BNST). Along these lines, Hermans and colleagues (Hermans et al., 2011) described greater salience-network connectivity during periods of anxiety associated with watching an aversive movie. In a previous study, we investigated network interactions when participants were in either threat (unpredictable mild shocks could be administered) or safe (no shocks possible) blocks (McMenamin et al., 2014). We found that the salience network exhibited a transient increase (following block onset) in network cohesion (that is, within-network functional connections increased) followed by decreased cohesion during a subsequent sustained period. Our study thus revealed changes to network organization during transient and sustained periods of threat.

Second, anxious anticipation is inherently temporal. Although previous studies have investigated how brain responses are sensitive to threat proximity (Mobbs et al., 2010, Somerville et al., 2010, Grupe et al., 2013), almost nothing is known about how patterns of brain co-activation (thus networks) *change* during dynamic manipulations of threat. For instance, although in our previous study (McMenamin et al., 2014) we characterized temporal aspects of network organization, threat itself was essentially constant (blocked).

To address these important gaps in the literature, we modulated threat dynamically during functional MRI scanning. Two circles moved on the screen for periods of 60 seconds, sometimes moving closer and sometimes moving apart (Fig. 1). If they touched, an unpleasant shock was delivered to the participant. We sought to determine how network properties changed during periods of approach (circles moving closer) and periods of retreat (circles moving apart). As in our previous study (McMenamin et al., 2014), we studied a set of regions spanning the salience, executive, and task-negative networks, given their involvement in cognitive and emotional processing (Yeo et al., 2011). In addition, we investigated a number of subcortical regions, many of which feature prominently in studies of the emotional brain (Pessoa, in press).

**Figure 1.**
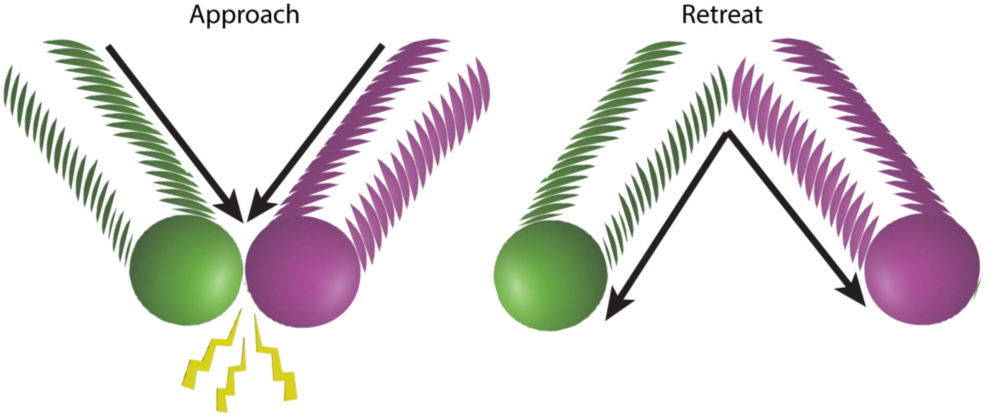
Stimuli paradigm. To create anxious states, over a period of 60 seconds, two circles with different colors moved around on the screen with some degree of randomness. When they collided with each other, an unpleasant mild electric shock was delivered.

We investigated the questions of interest within the framework of intersubject correlation analysis (Hasson et al., 2004). In this framework, time series data from voxels or regions of interest (ROI) are correlated *across* participants to determine “interpersonal synchronization” (Fig. 2A). Intersubject correlation is believed to reflect synchronization of mental states that are not simply explained by common evoked responses to perceptual features or cognitive demands (Nummenmaa et al., 2012, Lahnakoski et al., 2014, Nummenmaa et al., 2014, Ames et al., 2015).

**Figure 2.**
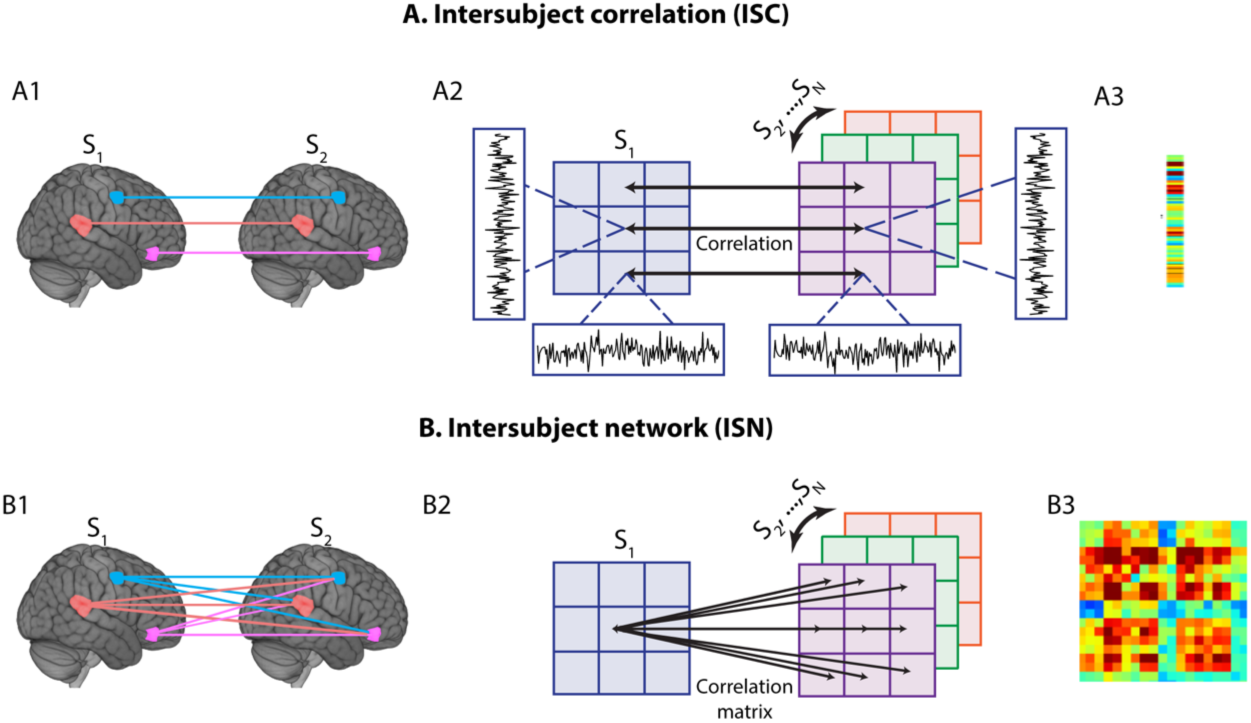
Intersubject correlation (ISC) and network analysis (ISN). A1) In ISC the correlation between the same region across different participants’ brains is calculated. **A2)** To calculate ISC, for each voxel or region of interest (ROI), the time series “left out” subject (S_1_) extracted, and correlated with the average time series of all other subjects (S_2_,…S_N_), for the same voxel/ROI. **A3)** To calculate the group level ISC, the results from A2 across all subjects are averaged. This creates a vector that contains the correlation of every voxel with itself (across participants). **B1)** In ISN the correlation between all pairs of ROIs across different brains is calculated. **B2)** To calculate ISN, for each voxel or ROI, the time series of a “left out” subjected (S_1_) is extracted and its correlation with the average time series across other subjects (S_2_,…,S_N_) is calculated. **B3)** The group level ISN is a #ROIs x #ROIs matrix, which shows the average of ISN from all subjects. Note that the vector in A3 corresponds to the diagonal of the matrix in B3, illustrating that intersubject networks provide a richer characterization of time series relationships.

Overall, our approach allowed us to test several questions about the brain basis of anxious anticipation. 1) How does dynamic threat reorganize the functional organization of large-scale brain networks? 2) How do network properties evolve during periods of threat approach and retreat? In particular, we sought to test the hypothesis that network organization evolves temporally during threat processing and that, for instance, salience-network cohesion increases/decreases with threat approach/retreat (Pessoa and McMenamin, 2016). 3) What is the relationship between cortical and subcortical regions important for threat processing during dynamic threat? 4) How are the amygdala and BNST involved in anxious anticipation? This question is important because, in particular, it is unclear if/how the amygdala is involved in conditions involving relatively prolonged and uncertain threat periods. The role of the BNST also remains unclear, as some have advocated that threat unpredictability is a critical determinant of its involvement (Alvarez et al., 2011).

## Materials and Methods

### Participants

Eighty-five participants (41 females, ages 18-40 years; average: 22.62, STD: 4.85) with normal or corrected-to-normal vision and no reported neurological or psychiatric disease were recruited from the University of Maryland community (of the original sample of 93, data from 7 subjects were discarded due to technical issues during data transfer [specifically, field maps were lost] and 1 other subject was removed because of poor structural-functional alignment). The project was approved by the University of Maryland College Park Institutional Review Board and all participants provided written informed consent before participation.

### Procedure and Stimuli

To create anxious states, two circles with different colors moved around on the screen randomly. When they collided with each other, an unpleasant mild electric shock was delivered. Overall, the proximity and relative velocity of the circles were used to influence the threat state. The position of each circle (on the plane), ***x***_t_, was defined based on its previous position, ***x***_*t*−1_, plus a random displacement, Δ***x***_*t*_:

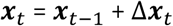

The magnitude and direction of the displacement was calculated by combining a normal random distribution with a momentum term to ensure motion smoothness, while at the same time remaining (relatively) unpredictable to the participants. Specifically, the displacement was updated every 50 ms as follows:

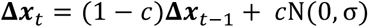

where *c = 0.2* and N(0, σ) indicates the normal distribution with 0 mean and standard deviation 1. The position and amount of displacement of each circle was updated independently.

Visual stimuli were presented using PsychoPy (http://www.psychopy.org/) and viewed on a projection screen via a mirror mounted to the scanner’s head coil. The total experiment included 6 runs, each of which had 6 blocks (3/85 participants had only 5 runs). In each block, the circles appeared on the screen and moved around for 60 seconds; blocks were preceded by a 15-second blank screen. Each run ended with 7 seconds of a blank screen.

To ensure that the effects of threat proximity and approach were uncorrelated, half of the blocks in each run were the temporally reversed versions of the other blocks in that run. Temporally reversing the stimulus trajectories guarantees that that proximity and approach are uncorrelated because reversing time changes the sign of the approach effect (that is, approach becomes retreat).

In each of the 6 runs the circles collided a total of 8 times in 4 out of the 6 blocks (3 shocks maximum per block); each collision resulted in the delivery of an electric shock. The 500-ms electric shock was delivered by an electric stimulator (Coulbourn Instruments, PA, USA) to the fourth and fifth fingers of the non-dominant left hand via MRI-compatible electrodes. To calibrate the intensity of the shock, each participant was asked to choose his/her own stimulation level immediately prior to functional imaging, such that the stimulus would be “highly unpleasant but not painful.” After each run, participants were asked about the unpleasantness of the stimulus in order to re-calibrate shock strength, if needed.

To optimize the experimental design, 10,000 candidate stimuli trajectories and block orders were generated. We then selected 6 runs which minimized collinearity between all predictors of interest (see below), measured as the sum of respective variance inflation factors (Neter et al., 1996).

### Stimulus Conditions

We defined two conditions, “approach” and “retreat,” based on whether the circles were moving toward or away from each other. Time points were only considered for analysis if the Euclidian distance between the circles was at least 75% of the maximum distance that the circles would exhibit during the whole experiment; otherwise, the data were not employed in the analysis. The rationale behind this was that, when the circles were far from each other, participants reported that they did not really pay as much attention to them. Therefore, we reasoned that the analysis should focus on the time points during which the circles were in (relative) closer proximity to each other. Investigation of the exploratory set revealed that the particular cutoff was not critical for the effects investigated (in the exploratory set only) and that values at least between 65% and 85% were adequate; based on the exploratory set results we chose a cutoff value of 75%.

### MRI data acquisition

Functional and structural MRI data were acquired using a 3T Siemens TRIO scanner with a 32-channel head coil. First, a high-resolution T2-weighted anatomical scan using Siemens’s SPACE sequence (0.8 mm isotropic) was collected. Subsequently, we collected 457 functional EPI volumes using a multiband scanning sequence (Feinberg et al., 2010) with TR = 1.0 sec, TE = 39 ms, FOV = 210 mm, and multiband factor = 6. Each volume contained 66 non-overlapping oblique slices oriented 30° clockwise relative to the AC-PC axis (2.2 mm isotropic). In addition, a high-resolution T1-weighted MPRAGE anatomical scan (0.8 mm isotropic) was collected. Finally, double-echo field maps (TE1 = 4.92 ms, TE2 = 7.38 ms) were acquired with acquisition parameters matched to the functional data.

### Functional MRI preprocessing

To preprocess the functional and anatomical MRI data, a combination of packages and in-house scripts were used. The first three volumes of each functional run were discarded to account for equilibration effects. Slice-timing correction (with AFNI’s 3dTshift) used Fourier interpolation to align the onset times of every slice in a volume to the first acquisition slice, and then a six-parameter rigid body transformation (with AFNI’s 3dvolreg) corrected head motion within and between runs by spatially registering each volume to the first volume.

Skull stripping determines which voxels are to be considered part of the brain and, although conceptually simple, plays a very important role in successful subsequent co-registration and normalization steps. Currently, available packages perform sub-optimally in specific cases, and mistakes in the brain-to-skull segmentation can be easily identified. Accordingly, to skull strip the T1 high-resolution anatomical image (which was rotated to match the oblique plane of the functional data with AFNI’s 3dWarp), we employed six different packages (ANTs (Avants et al., 2011), AFNI (Cox, 1996; http://afni.nimh.nih.gov/): http://afni.nimh.nih.gov/, ROBEX (Iglesias et al., 2011): https://www.nitrc.org/projects/robex), FSL: http://fsl.fmrib.ox.ac.uk/fsl/fslwiki/, SPM: http://www.fil.ion.ucl.ac.uk/spm/, and Brainsuite (Shattuck and Leahy, 2002) and employed a “voting scheme” as follows: based on T1 data, a voxel was considered to be part of the brain if 4/6 packages estimated it to be a brain voxel; otherwise the voxel was not considered to be brain tissue (for 6 subjects whose T1 data were lost due to issues during data transfer, the T2 image was used instead and only the ANTs package was used for skull-stripping).

Subsequently, FSL was used to process field map images and create a phase-distortion map for each participant (bet and fsl_prepare_fieldmap). FSL’s epi_reg was then used to apply boundary-based co-registration to align the unwarped mean volume registered EPI images with the skull-stripped anatomical image (T1 or T2) along with simultaneous EPI distortion-correction (Greve and Fischl, 2009).

Next, ANTS was used to determine a nonlinear transformation that mapped the skull-stripped anatomical image (T1 or T2) to the MNI152 template (interpolated to 1-mm isotropic voxels). Finally, ANTS combined the nonlinear transformations from co-registration/unwarping (from mapping mean functional EPI images to the anatomical T1 or T2) and normalization (from mapping T1 or T2 to the MNI template) into a single transformation that was applied to map registered functional volumes of functional data to standard space (interpolated to 2-mm isotropic voxels). In this process, ANTS also utilized the field maps to simultaneously minimize EPI distortion.

### Time series data

As we sought to characterize patterns of co-activation, time series data were initially processed so as to remove the estimated contributions of the paradigm. To do so, we ran multiple linear regression (with AFNI’s 3dDeconvolve) on the preprocessed functional data with the goal of estimating the residual time series after the inclusion of the following regressors: proximity, velocity, velocity x proximity, and visual motion. The regressors were determined based on the circle positions on the screen. Proximity was defined as the Euclidean distance between the two circles. Velocity was the discrete temporal difference of proximity. The visual motion regressor was defined as the velocity tangential to the difference vector of the combined circle-to-circle stimulus, and was calculated by multiplying the angular velocity of the difference vector by the proximity (and accounted for motion energy orthogonal to the relative motion between the circles).

For each run, the regressors were obtained by first decimating the 20 Hz sample rate of stimuli information (used to compute circle paths) to the TR sample rate (1 Hz). Within each run, the regressors were mean-centered to reduce collinearity between simple effects (proximity, velocity) and the velocity x proximity interaction term, and convolved with a standard hemodynamic response based on the gamma-variate model (Cohen, 1997). In addition, we regressed out any potential block-sustained activation by including a regressor representing the blocks (60-second duration), which was convolved with the standard hemodynamic response. Other regressors included in the model comprised 6 motion parameters (3 linear displacements and 3 angular rotations), and their discrete temporal derivatives. Additionally, to model baseline and drifts of the MR signal, regressors corresponding to polynomial terms up to 4^th^ order were included (for each run separately). To minimize the shock effect, data points in a 15-sec window after shock delivery were discarded from all analyses. Finally, the residual time series for each run were z-scored separately.

The residual time series as defined above was used for the intersubject network analysis, whose main goal was to characterize networks during approach and retreat conditions, and contrast them. As specified above, approach and retreat varied dynamically throughout the blocks. Therefore, we employed a windowing procedure to extract data segments corresponding to the conditions of interest. Intuitively, the windowing allowed us to select segments of the time series associated with each condition and concatenate them across all runs, generating a final concatenated times series for each condition. First, for each block, the first 15 time points (15 seconds) were discarded, to minimize contributions from block onset. Segment type was determined by considering the velocity regressors (which determined approach vs. retreat). Specifically, transitions in the sign of the velocity regressor indicated the start of a segment type (approach or retreat), to which a 5-second lag was added to account for hemodynamics. Furthermore, based on the 75% cutoff described earlier, data were discarded based on proximity data; in other words, we only considered data in the “75% near space.” Finally, all the time points across all blocks and runs assigned to each condition were concatenated for that condition, and constituted the time series data for the condition.

### Exploratory and test sets

The total dataset was subdivided into “exploratory” and “test” sets. The idea was to use the exploratory set to fix specific processing choices; with the entire processing pipeline fixed, statistical testing was then applied to separate data in the test set. The size of the exploratory set (N=37) was determined arbitrarily and based on splitting the data available at a certain date during the data acquisition process; as scanning continued for a bit longer, the test set contained a larger number of participants (N=48; the original goal being to have approximately 50 participants). To reiterate, the results reported here are based on the test set alone; thus, processing choices were not optimized or tuned to the test sample, by design.

### Regions of interest (ROIs)

We investigated three networks widely studied in the literature: salience, executive, and task negative. From these networks, we employed cortical ROIs (defined as 5-mm radius spheres) based on the center coordinates provided by previous studies: salience network (Hermans et al., 2011) (13 regions), executive network (Seeley et al., 2007) (12 regions), and task-negative network (Fox et al., 2005) (12 regions).

Based on our goal of investigating threat/anxiety, we included additional subcortical ROIs: amygdala, hippocampus, cerebellum, PAG, habenula, and BNST. For the amygdala, we considered the subregions defined by the Nacewicz et al. (Nacewicz B.M. et al., 2014) parcellation, specifically: lateral amygdala; basolateral/medial amygdala; cortical nucleus plus amygdalo-hippocampal area; central plus medial nuclei. For the hippocampus, we focused on its anterior portion because the rodent literature has implicated the ventral hippocampus (believed to correspond to the anterior part in humans) in anxiety-related processing (Bannerman et al., 2004). The hippocampus ROI was defined by using the hippocampus mask from FreeSurfer and cutting it at the y = +21 plane (MNI coordinates). For the cerebellum, a meta-analysis indicated that Lobules I-IV and Crus II were involved in emotion-related processing (Riedel et al., 2015). Masks for these regions were obtained from the cerebellum parcellation available in FSL (Diedrichsen et al., 2009) (called VIIa Crus II region in the FSL atlas). For the PAG, we modified the mask by (Roy et al., 2014), which was dilated by 1 voxel; in addition, we manually removed the voxels that extended above/below the superior/inferior limits of the original ROI, and those overlapping cerebrospinal fluid. The habenula has been implicated in emotional/motivational processing (Hikosaka, 2010), and here we employed a mask defined according to the Morel atlas, as defined in (Krauth et al., 2010). For the BNST, we employed a recently developed mask based on 7 Tesla data but defined having in mind 3 Tesla data (Theiss et al., 2017).

If two cortical or subcortical ROIs abutted each other, each mask was eroded by 1 voxel from the touching boundary to minimize any potential data “spill over.” The exceptions to this rule were the amygdala subregions, BNST, and habenula, because of their very small volume.

Based on exploratory set results, for test set inferences, we focused on the following subcortical ROIs: right lateral amygdala; right PAG, right habenula, left cerebellum crus and right BNST. For completeness, and to foster cross-study continuity, we include in Supplementary Figs. 1-3 results from all regions investigated in the exploratory set.

### Intersubject functional network

In previous intersubject studies, two approaches have been used. First, the time series of an ROI (or voxel) in one subject is correlated with time series data of the same ROI (or voxel) in the remaining subjects (for example, it can be correlated with the average ROI time series data across the “remaining” subjects), yielding a resulting intersubject correlation map (Hasson et al., 2004). Second, intersubject seed-based analysis can be performed, in which the time series of an ROI in one subject is correlated with time series data of a group of ROIs (or voxels) of other subjects (Simony et al., 2016).

A simple, yet powerful extension is to consider intersubject correlations across all pairs of ROIs, which allows the application of the technique to networks. The procedure to generate an intersubject network is specified in Fig. 7. For a given ROI, first a subject’s s data is held out (***y***_s_), and the rest of the subjects’ time series is averaged (***ӯ***_−*s*_). Then, the Pearson correlation between the left-out data and the corresponding data in the remaining subjects is computed: corr(***ӯ**_*s*_,**y**_–s_*). This basic operation is repeated for all pairs of ROIs to compute an intersubject network for the held-out subject (**ISN**_S_). (For compactness, all of the ROI time series are stacked into a matrix *Y* in the algorithm of Fig. 7.) Thus, the **ISN**_S_ is an ***N×N*** matrix, where *N* is the number of ROIs, and the *ij-th* matrix element contains the correlation coefficient between the *i*-th ROI time series of the held-out subject and the average of time series of the *j*-th ROI of the remaining subjects.

**Figure 3.**
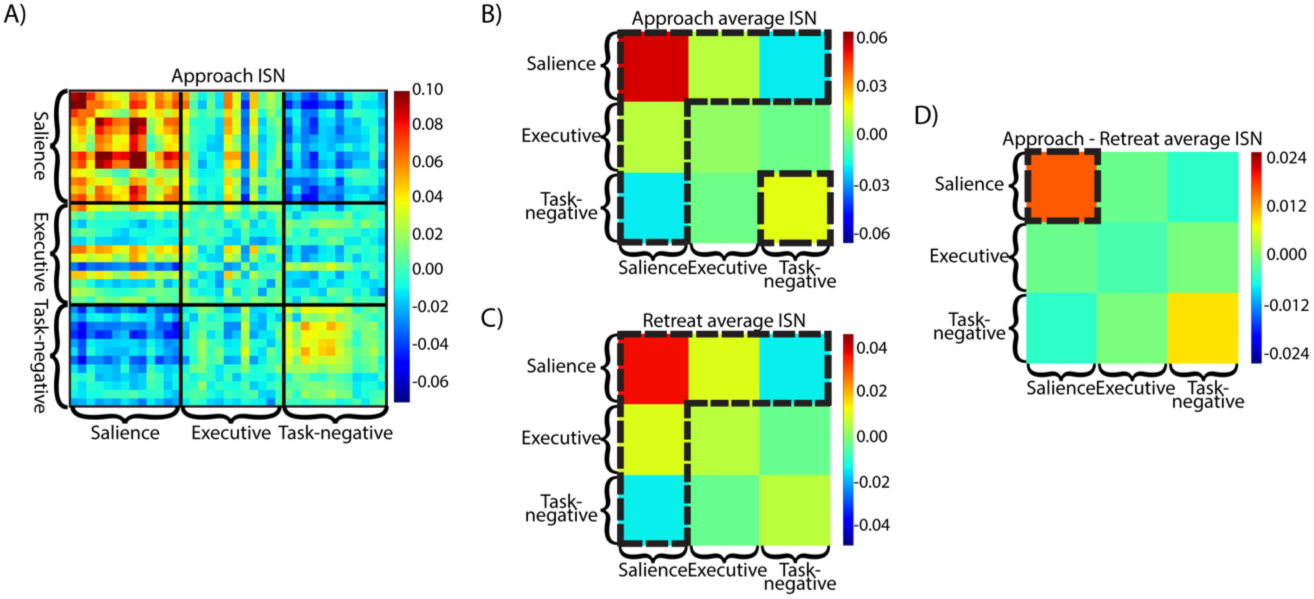
Intersubject group networks. **A)** Intersubject network (ISN) during threat approach periods. ROI order corresponds that that of Table 1. **B-D)** Average ISN values for each network at approach (B), retreat (C), and approach minus retreat (D). The dark rectangles surround all of the blocks with significant values (p < 0.01); note that each of the 9 blocks was treated as a unity(the outline extended over multiple of them for diagramming convenience). The color bars indicate differences in intersubject correlation.

**Table 1:**
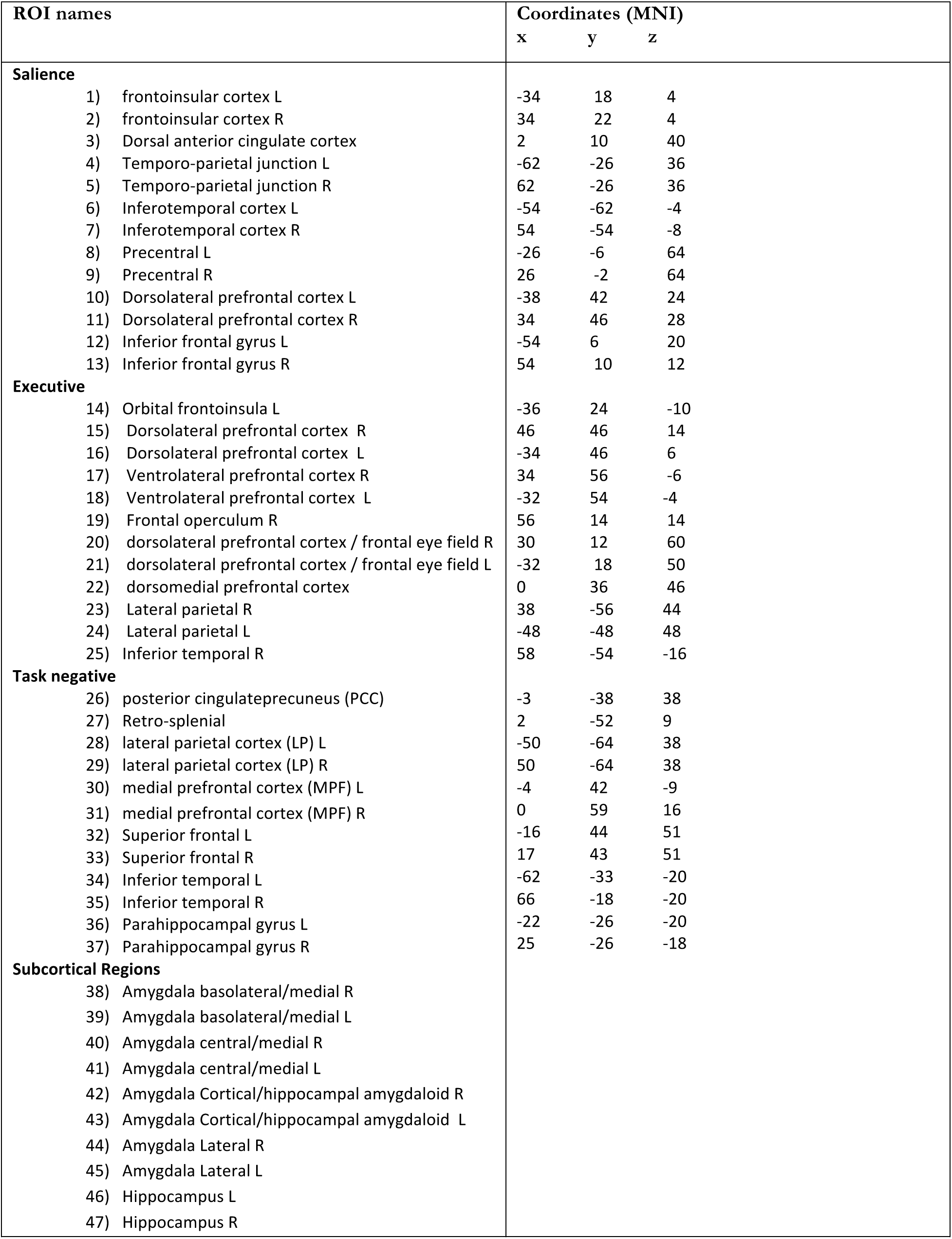
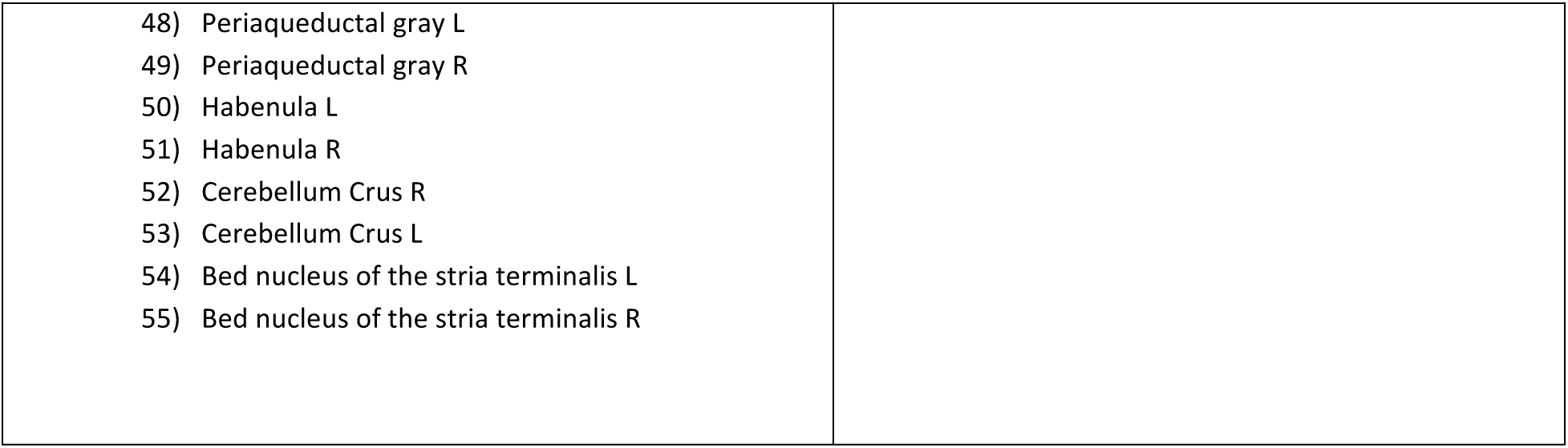
List of Cortical and Subcortical Regions of Interest (ROIs) Cortical ROIs were defined via 5-mm radius spheres centered on MNI coordinates provided below. Subcortical ROIs were defined anatomically (see text for details).

| ROI names | Coordinates (MNI) |  |  |
| --- | --- | --- | --- |
|  | x | y | z |
| **Salience** |  |  |  |
| 1) frontoinsula cortex L | -34 | 18 | 4 |
| 2) frontoinsula cortex R | 34 | 22 | 4 |
| 3) Dorsal anterior cingulate cortex | 2 | 10 | 40 |
| 4) Temporo-parietal junction L | -62 | -26 | 36 |
| 5) Temporo-parietal junction R | 62 | -26 | 36 |
| 6) Inferotemporal cortex L | -54 | -62 | -4 |
| 7) Inferotemporal cortex R | 54 | -54 | -8 |
| 8) Precentral L | -26 | -6 | 64 |
| 9) Precentral R | 26 | -2 | 64 |
| 10) Dorsolateral prefrontal cortex L | -38 | 42 | 24 |
| 11) Dorsolateral prefrontal cortex R | 34 | 46 | 28 |
| 12) Inferior frontal gyrus L | -54 | 6 | 20 |
| 13) Inferior frontal gyrus R | 54 | 10 | 12 |
| **Executive** |  |  |  |
| 14) Orbital frontoinsula L | -36 | 24 | -10 |
| 15) Dorsolateral prefrontal cortex R | 46 | 46 | 14 |
| 16) Dorsolateral prefrontal cortex L | -34 | 46 | 6 |
| 17) Ventrolateral prefrontal cortex R | 34 | 56 | -6 |
| 18) Ventrolateral prefrontal cortex L | -32 | 54 | -4 |
| 19) Frontal operculum R | 56 | 14 | 14 |
| 20) dorsolateral prefrontal cortex / frontal eye field R | 30 | 12 | 60 |
| 21) dorsolateral prefrontal cortex / frontal eye field L | -32 | 18 | 50 |
| 22) dorsomedial prefrontal cortex | 0 | 36 | 46 |
| 23) Lateral parietal R | 38 | -56 | 44 |
| 24) Lateral parietal L | -48 | -48 | 48 |
| 25) Inferior temporal R | 58 | -54 | -16 |
| **Task negative** |  |  |  |
| 26) posterior cingulateprecuneus (PCC) | -3 | -38 | 38 |
| 27) Retro-splenial | 2 | -52 | 9 |
| 28) lateral parietal cortex (LP) L | -50 | -64 | 38 |
| 29) lateral parietal cortex (LP) R | 50 | -64 | 38 |
| 30) medial prefrontal cortex (MPF) L | -4 | 42 | -9 |
| 31) medial prefrontal cortex (MPF) R | 0 | 59 | 16 |
| 32) Superior frontal L | -16 | 44 | 51 |
| 33) Superior frontal R | 17 | 43 | 51 |
| 34) Inferior temporal L | -62 | -33 | -20 |
| 35) Inferior temporal R | 66 | -18 | -20 |
| 36) Parahippocampal gyrus L | -22 | -26 | -20 |
| 37) Parahippocampal gyrus R | 25 | -26 | -18 |
| **Subcortical Regions** |  |  |  |
| 38) Amygdala basolateral/medial R |  |  |  |
| 39) Amygdala basolateral/medial L |  |  |  |
| 40) Amygdala central/medial R |  |  |  |
| 41) Amygdala central/medial L |  |  |  |
| 42) Amygdala Cortical/hippocampal amygdaloid R |  |  |  |
| 43) Amygdala Cortical/hippocampal amygdaloid L |  |  |  |
| 44) Amygdala Lateral R |  |  |  |
| 45) Amygdala Lateral L |  |  |  |
| 46) Hippocampus L |  |  |  |
| 47) Hippocampus R |  |  |  |

|  |
| --- |
| 48) Periaqueductal gray L |
| 49) Periaqueductal gray R |
| 50) Habenula L |
| 51) Habenula R |
| 52) Cerebellum Crus R |
| 53) Cerebellum Crus L |
| 54) Bed nucleus of the stria terminalis L |
| 55) Bed nucleus of the stria terminalis R |

**Figure 4.**
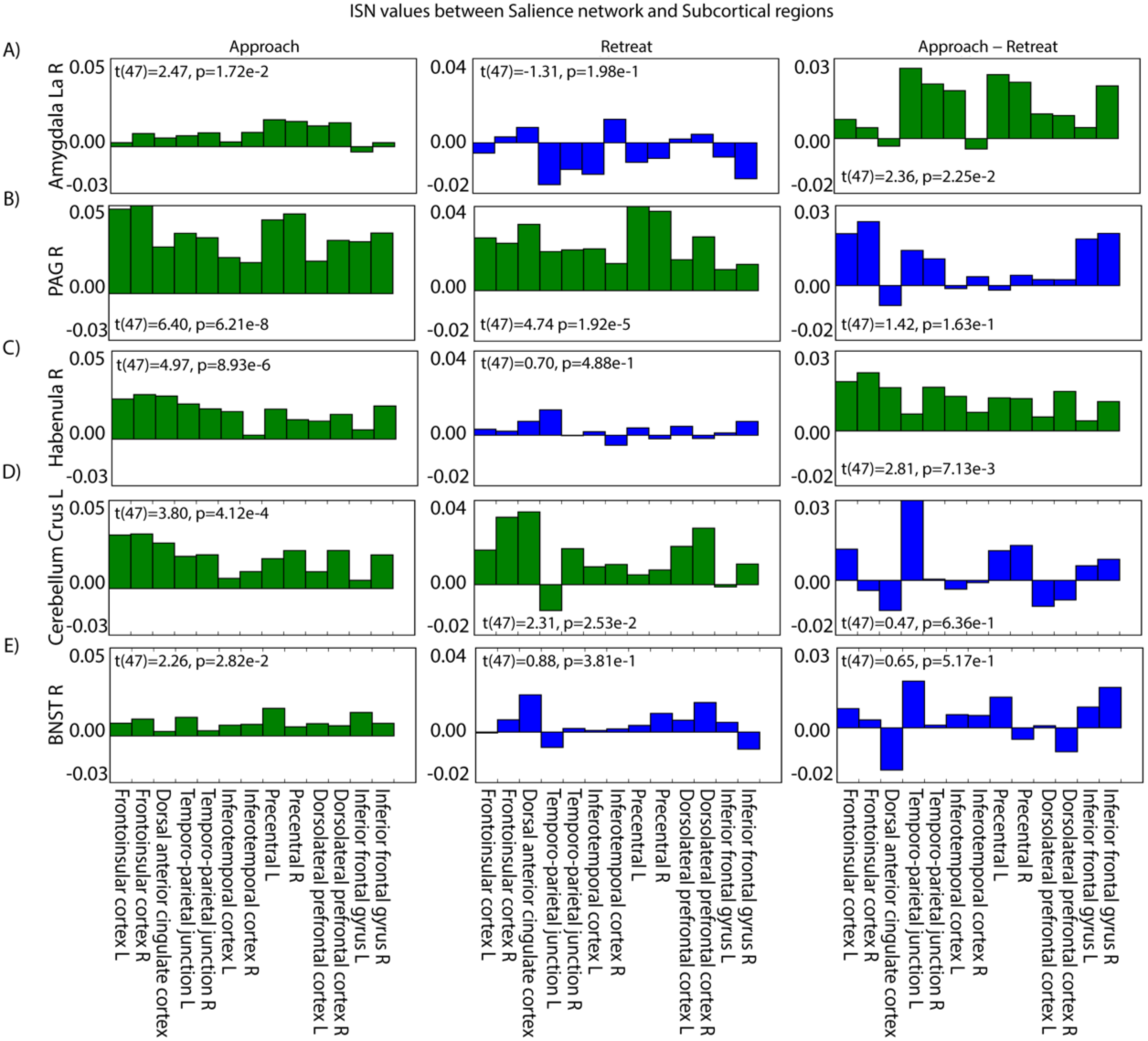
Intersubject functional connections between the salience network and subcortical regions. **A)** Amygdala La R; **B)** PAG R; **C)** Habenula R; **D)** Cerebellum Crus L. **E)** BNST R. L/R: left, right; Amygdala La: lateral amygdala, PAG: periaqueductal gray, BNST: Bed nucleus of the stria terminalis. For statistical tests, the entire subcortical region was treated as a unit and cohesion between the region and the salience network was tested. Regions shown with green bar plots were significant (p<0.05).

**Figure 5.**
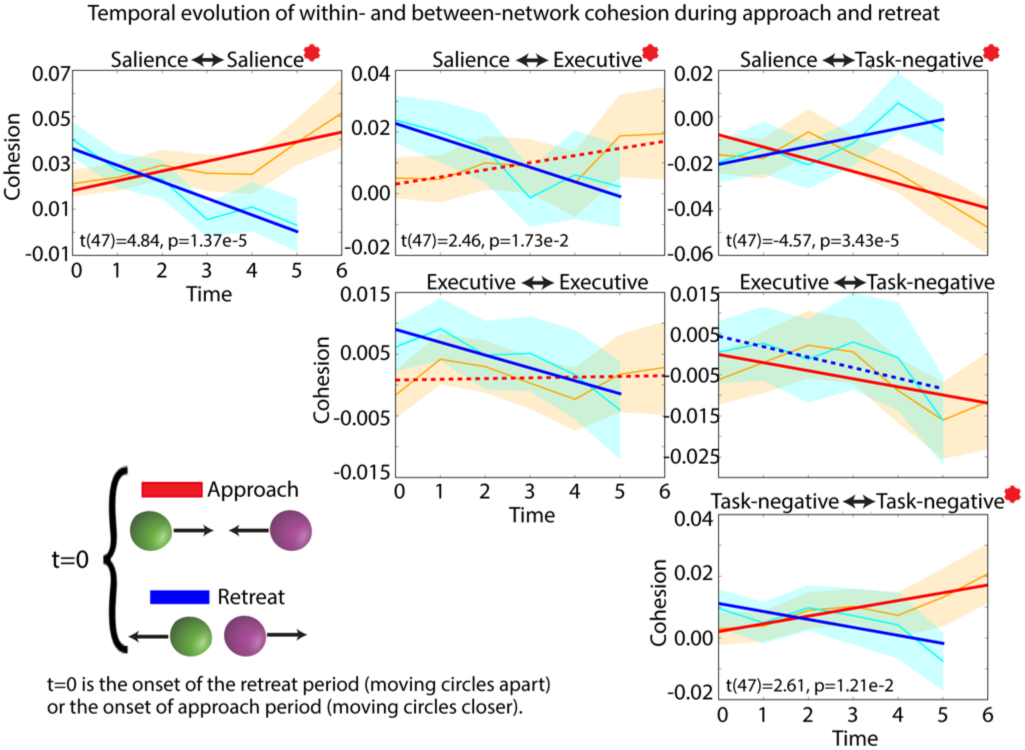
Temporal evolution of cohesion during approach and retreat segments. Within- and between-network cohesion during approach and retreat for the salience, executive, and task-negative networks. For example, as the circles approach each other, the cohesion within the salience network increased; when the circles retreated, the cohesion within the salience network gradually decreased. The orange line shows cohesion values for approach at different times (with a 90% confidence band); the cyan line shows cohesion values for retreat at different times (90% confidence band). The red/blue line shows the least-squares linear fit to cohesion values during approach/retreat; solid lines indicate fits that were statistically significant (p < 0.05). Time is in seconds (TR = 1 sec); the y-axis shows cohesion (summed degree). The red star indicates that the slope difference was statistically significant (statistical values provided at the bottom left)

**Figure 6.**
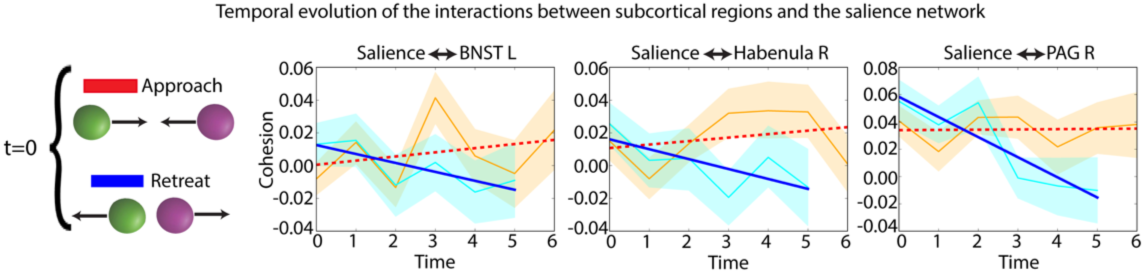
Temporal evolution of cohesion between subcortical regions and the salience network. Although cohesion did not increase robustly during approach periods, cohesion decreased as a function of time during retreat for the left BNST, right habenula, and right PAG. Conventions as in Fig. 5.

**Figure 7.**
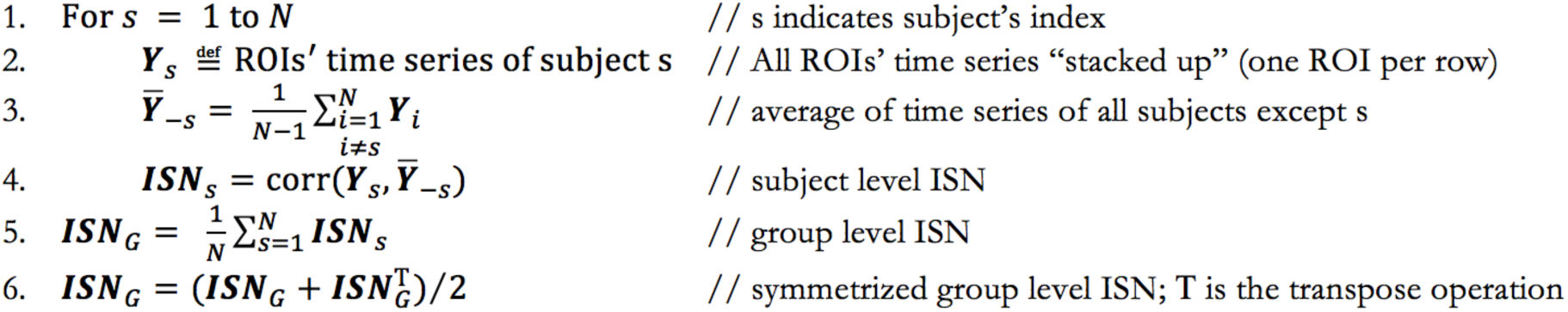
Computation of group intersubject network. The operator corr corresponds to Pearson correlation.

This procedure is then repeated for all subjects. A group matrix (**ISN**_G_) is then obtained by averaging across all subjects. Note that the resulting intersubject network is not necessarily symmetric, because, for each ROI, the time series in the held-out subject (***y**_s_*) is not necessarily equal to average of all other subjects’ time series (***ӯ***_–*s*_). While the **ISN**_G_ matrix in practice will be near symmetric, a simple and intuitive way to mathematically accomplish symmetry is to average the group-level intersubject network with its transpose (where row and column indexes are flipped), leading to a final symmetric group matrix. Finally, the procedures above were performed, separately, for the approach and retreat conditions generating the matrices **ISN**_G,APPROACH_ and **ISN**_G_,_RETREAT_. Finally, note that in our method the diagonal is also computed because data at a given diagonal entry {*i*,*i*} is computed across different brains.

### Dynamic intersubject networks

Considering all data points simultaneously, as done thus far, gives us a static understanding of intersubject networks. Here, we sought to investigate dynamic aspects of network organization. To do so, we computed intersubject networks at each time *t* and investigated how network properties changed across segments of approach and retreat. Thus, for each segment type (approach and retreat, separately), we considered all of the ***t* = 0** time points, then ***t* = 1** time points, and so on, separately. The goal was to generate a time series of data at ***t* = 0** by concatenating all of the ***t* = 0** data across segments. To do so, we concatenated the ***t* =** ***k*** points (for a fixed *k*) across segments (Fig. 8). To account for the hemodynamic delay, we discarded the first 5 seconds of each segment. In this manner, we determined ISN_t_=_0_, ISN_t=i_, and so on. We considered intersubject networks for at ***t* = 0, ⋯ ,6** for the approach condition (circles moving closer to each other) and ***t* = 0, ⋯ ,5** for the retreat condition (circles are moving away from each other). This was done such that at least 20 data points were available per condition and time “slices” (note that less data was available for the retreat condition, as some points were discarded following shock events).

**Figure 8.**
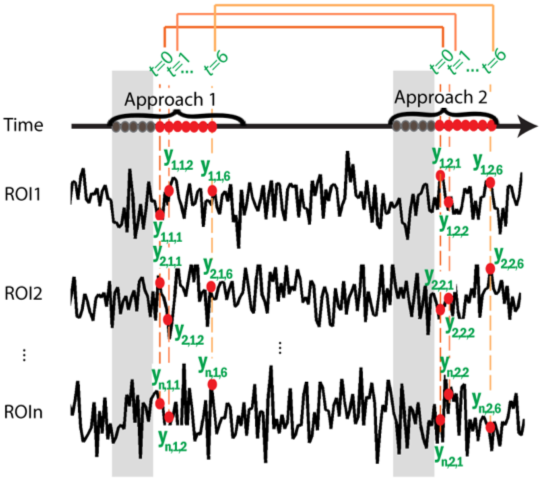
Procedure for data concatenation used in evaluating intersubject network dynamics. Procedure for data concatenation used in evaluating intersubject network dynamics. Data point *y* for each time point *t* was used to compute intersubject networks for approach and retreat, separately. Data point indexes: ROI#, approach/retreat segment# (two approach data segments are illustrated), and time within segment. Briefly, time was used to “slice” and “concatenate” through the ROI time series. Thus, we generated a data time series at *t*=0 by concatenating all of the ***t*=0** sample (across same-conditionsegments across blocks and runs), did the same for *t*=1, and so on. To account for the hemodynamic delay, we discarded the first 5 seconds of each segment (gray part). The resulting data per ROI, time point, and condition, was then investigated in terms of dynamic properties. Time series data were simulated for illustration.

### Within- and between-network cohesion

To measure the strength/cohesion of connections within and between networks, we utilized within- and between-network degree, respectively. The cohesion between network
***N_1_*** and network ***N*_2_**, ***C*_*N*_1__, *N*_2_**, was defined as follows:

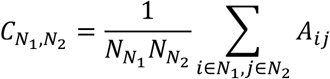
where ***N*_*N*1_** is the number of ROIs in network ***N***_1_, and ***N**_**N**_*_2_ is the number of ROIs in network ***N***_2_. ***A*_*ij*_** is the **ISN**_G_ value between i-th ROI and j-th ROI when i-th ROI belongs to network ***N**_1_* and j-th ROI belongs to network ***N***_2_. If the above formula, ***N**_1_* and ***N***_2_ are the same network, then above formula calculates within-network cohesion. In this case both i-th and j-th ROIs belong to the same network.

Defining network cohesion in terms of degree had two advantages. First, we considered both positive and negative weights, unlike most approaches that discard negative weights because many network measures (such as *efficiency*) do not easily handle negative values (Newman, 2010b). Most network measures also do not handle self-connections (Newman, 2010a), which in standard analysis are not informative anyway (***A_ii_ =1***). Here, we considered functional connections between the same region (across brains), which could be incorporated in within-network cohesion by considering the terms *A_ii_* in the computation of cohesion, *C*.

To evaluate functional connections between subcortical regions and the salience network (Fig. 4), we computed a cohesion index that summed all functional connections between a specific subcortical region and all nodes of the salience network. This was performed for the approach and retreat conditions, separately. To test for cohesion, a one-sample t-test (against zero) was employed. A paired t-test was employed to compare cohesion between approach to retreat conditions.

To study dynamic changes to network cohesion, ISN_G_ was computed as outlined above at each time *t* (Fig. 5). To test for time effects, linear regression was employed, and applied separately to the approach and retreat conditions.

## Results

### Intersubject network analysis and statistical approach

Standard network analysis of functional MRI data is based on a adjacency matrix in which each entry is the correlation between time series data for two ROIs within the same participant (Bullmore and Sporns, 2009). Here, we employ a method to extend intersubject correlation analysis to networks. Although we developed this method independently (Najafi and Pessoa, 2016),Simony and colleagues developed essentially the same method and applied it to the study of understanding the task-negative network during narrative comprehension (Simony et al., 2016). We call our version of the method *intersubject network analysis*. Specifically, the correlations across all pairs of regions is computed (Fig. 2B), generating a correlation matrix that can be investigated via graph theory techniques (Newman, 2010a).

By correlating time series data across participants, intersubject network analysis captures temporal signal properties that are shared by them, deemphasizing fluctuations that are incidental and observed in individual participants (Simony et al., 2016). Another important property of intersubject network analysis is that it can consider the correlation of a region with itself. Whereas in standard analysis this is uninformative (a region’s correlation with itself is by definition 1), in intersubject network analysis the correlation is meaningful (and informative) because the time series data come from different brains.

To develop the method and its application to the dynamic threat paradigm without “peeking” into data, we applied it first to a subset of our entire dataset, which we call the “exploratory set” (N=37), and was used to fix particular processing choices. The results described here were obtained in a separate “test set” (N=48) independent from the exploratory set. Our goal was to enhance reproducibility in a research area plagued by the “curse of flexibility.” For example, Poldrack and colleagues (Poldrack et al., 2017) recently enumerated 69,120 different workflows for basic functional MRI analysis alone. We advocate the present approach with exploratory and test sets to imaging studies that do not target very specific questions (which we believe are rarer than typically acknowledged), and/or that include novel methodology (as in the present case). Note that, although we refer to our sets as “exploratory” and “test,” our goal was not to attempt discovery and replication in a single study, as in genomics, for example. Specifically, the objective of using an exploratory set was to develop the method and to narrow down the brain regions being investigated.

The total experiment included 6 runs, each of which included 6 blocks. In each block, the circles moved on the screen for 60 seconds; blocks were separated by a 15-second blank screen. We investigated functional connectivity of several regions that are challenging to image with functional MRI, including amygdala subnuclei, the BNST, and the habenula. Although great care was taken at co-registration and functional data were not smoothed, we suggest that region labels be considered “putative” insofar as higher functional resolution would be required for clearer anatomical attribution. See Methods for further information.

### Network cohesion

We determined intersubject correlation matrices for the approach and retreat conditions (see Methods), which allowed us to determine within- and between network cohesion (based on node *degree*) for the two conditions, and to compare cohesion values during approach vs. retreat. Positive cohesion values indicate that correlations between regions within a network or between networks were on average positive; negative cohesion values indicate that they were on average negative. Note that although our measure of cohesion was based on degree, it is not subject to the recent criticism of using degree when estimating node importance (Power et al., 2013), because that was not our goal here. Importantly, by utilizing node degree, we could parsimoniously employ both positive and negative weights, thus providing a better characterization of network “strength.”

During approach (Fig. 3A), positive network cohesion was detected within the salience network (one-sample *t* test; t(47)= 8.91; p=9.45e-12), and between the salience and executive networks (one-sample *t* test; t(47)=2.8; p=5.92e-3); negative cohesion was detected between the salience and task-negative networks (one-sample *t* test; t(*47*)=-5.61; p=9.68e-7) (nodes tended to be negatively correlated). Furthermore, within-network cohesion was positive in the task-negative network (one-sample *t* test; t(*47*)=3.42; p=1.26e-3) (Fig. 3B-D plots the results as summary “block matrices” for convenience). Interestingly, a similar pattern of results was observed during retreat (in all cases one-sample t tests; within salience: t(47)=6.63; p=2.74e-8; between salience and executive: t(*47*)=3.14; p=2.89e-3; between salience and task-negative: t(*47*)=-3.61; p=7.17e-4). Finally, the direct comparison between approach vs. retreat revealed increased cohesion in the salience network during approach vs. retreat (two-sample paired *t* test; t(*47*)=2.67; p=1.01e-2).

In brief, during threat processing (including both approach and retreat periods) both the salience network and its interactions with other networks exhibited the most conspicuous patterns of correlation. In addition, the task-negative network also exhibited positive cohesion during approach.

### Subcortical regions

Several subcortical regions are involved in threat processing. Furthermore, it has been suggested that some subcortical regions are functionally linked with the salience network under threat conditions (Hermans et al., 2011). Based on existing literature, we considered the amygdala (subdivided into subregions), anterior hippocampus, periaqueductal gray (PAG), and BNST. Based on analysis using the exploratory set, we also report test-set results on the habenula and the cerebellum crus.

Fig. 4 displays intersubject functional correlations between the salience network and subcortical regions. To evaluate functional connections between subcortical regions and the salience network, we computed a cohesion index that treated the subcortical region as a unit (thus summing degree across all nodes of the salience network). For each of reference, statistical values are provided for each subcortical region and condition in Fig. 4 (see Methods). During approach, several subcortical regions were positively correlated with the salience network, including the right lateral amygdala, right/left PAG, right/left habenula, left Cerebellum crus, and right BNST. As the pattern was somewhat similar during retreat, only the following regions exhibited robust difference for approach relative to retreat: right basolateral/medial amygdala, right lateral amygdala, left PAG, and right habenula.

### Dynamics

As threat level was varied dynamically, we investigated how the intersubject correlation matrix evolved temporally. Fig. 5 shows the temporal evolution of within- and between-network cohesion during approach and retreat for the salience, executive, and task-negative networks. For the salience network, within-network cohesion increased during approach periods and decreased during retreat periods (for ease of reference, statistical values are provided in the figure). Notice that the reverse was observed for cohesion between the salience and task-negative networks. Overall, all networks exhibited dynamic changes during approach and/or retreat periods (that is, at least one of the slopes was statistically significant).

We also investigated the evolution of the interactions between subcortical regions and the salience network. Although cohesion did not increase robustly during approach periods, decreased cohesion was detected during retreat periods for the left BNST, right habenula, and right PAG (Fig. 6), revealing that their functional association with the salience network is dynamic.

## Discussion

In the present study, we employed intersubject network analysis to investigate the properties of large-scale networks during threat approach and retreat. A central aim was to investigate the evolution of network properties as threat level varied dynamically. Threat altered network cohesion across the salience, executive, and task-negative networks, as well as subcortical regions. Importantly, cohesion within and between networks changed dynamically as threat imminence increased and decreased (as circles moved closer and farther to each other). Next, we discuss the implications of our main findings.

Standard intersubject correlation analysis has been used to investigate “synchrony” across brains when participants watch the same movie or during other naturalistic conditions, such as hearing extended narratives (Hasson et al., 2004, Stephens et al., 2010, Nummenmaa et al.,2012, Nummenmaa et al., 2014). The original formulation was inherently bivariate and considered the same voxel (or region) across participants. The method was recently extended so that a specific voxel/region in one person could be correlated with multiple voxels/regions in other participants (Simony et al., 2016). Independently, we formulated essentially the same method (Najafi and Pessoa, 2016) to perform *intersubject network analysis* with the aim of understanding network organization during dynamic threat. Although we had a specific analysis goal in mind (evaluating network cohesion), more generally, the full range of techniques developed for network analysis can be applied to intersubject data.

Intersubject analyses in general have the advantage that they increase the signal-to-noise ratio by filtering out unwanted contributions to the measured BOLD signal (Simony et al.,2016). This is particularly important for head motion, which can induce significant within-participant correlations (Van Dijk et al., 2012). By computing correlations across participants, the approach essentially eliminates this issue (on the test dataset, head motion parameters exhibited a correlation across subjects of .02).

A central finding of our study was that cohesion within and between networks changed dynamically during periods of approach and retreat. This adds to findings from recent studies that showed how large-scale networks are reorganized during periods of threat (Hermans et al., 2011, McMenamin et al., 2014). Consistent with previous studies, the salience network cohesion increased for approach relative to retreat. Critically, cohesion was not static during approach/retreat segments, but dynamically increased during approach and decreased during retreat. The cohesion between the salience and executive networks followed the same pattern. Notably, the reverse was observed between the salience and task-negative networks. Thus, the salience and task-negative networks cohered more strongly as the circles moved away from each other; movement toward each other made the networks less cohesive. Overall, our findings demonstrate that network cohesion is a dynamic property that depends on threat proximity.

The salience network comprises multiple regions in parietal, frontal, and insular cortices (Seeley et al., 2007, Menon and Uddin, 2010). Sometimes subcortical regions are listed as part of the network, most notably the amygdala and PAG (Seeley et al., 2007). In the present study, we hypothesized that an extended set of subcortical regions would be closely associated functionally to the salience network. Indeed, this was observed in our data, including, during threat approach, the right lateral amygdala, right/left periaqueductal gray, right/left habenula, cerebellum crus, cerebellum lobes, and right BNST. The present study shows that, in the context of threat processing, these regions are functionally linked to salience processing in paradigms involving threat, and not only during the resting state (see also (Hermans et al., 2011). Importantly, it also shows that this property changes during periods of threat approach relative to retreat, such as in the right lateral amygdala (Fig. 4).

The findings about the amygdala are particularly noteworthy. Whereas the amygdala is engaged by emotion-laden stimuli and conditions involving acute threat (as in aversive conditioning paradigms), its involvement in potential threat (where threat is not proximal and is relatively uncertain) is less clear (Davis et al., 2010b). Some human neuroimaging studies even observed deactivations of the amygdala during conditions of potential threat (Pruessner et al., 2008, Wager et al., 2009, Choi et al., 2012). Here, we saw that multiple amygdala subregions exhibited increased differential functional correlation (approach greater than retreat) with the salience network. These findings are important, because they show that the amygdala is involved during some forms of potential threat (such as the one studied here), as revealed by changes in co-activation patterns (see also (Hermans et al., 2011, McMenamin et al., 2014). It also highlights the need to study functional connectivity and network properties, in addition to evoked responses.

The involvement of the BNST in potential threat was suggested in early work by Davis and colleagues (Davis and Shi, 1999) and has been investigated recently in rodent studies with new neurotechnologies (see (Tovote et al., 2015). Work in humans has revealed the involvement of the BNST in potential threat, too (for reviews, see (Fox et al., 2015, Shackman and Fox, 2016). The BNST is rather small and thus challenging to investigate in humans with functional MRI. Nevertheless, recent work at higher resolution and magnetic field strengths (such as 7 Tesla) has been used to generate anatomical masks (Avery et al., 2014, Torrisi et al., 2015), and these appear to be reasonable approximations even at the standard field strength of 3 Tesla (Theiss et al., 2017). An open question concerns the conditions leading to BNST engagement. While some studies suggest that uncertainty may be a major determinant of BNST responses (Alvarez et al., 2011), this is not entirely clear. For example, in a previous study, Mobbs and colleagues (Mobbs et al., 2010) found greater BNST responses for threat approach vs. retreat (although the authors only employed a single level of approach vs. retreat “level,” and the activation pattern was very diffuse, thus hard to attribute to the BNST with more confidence). In the present study, the right BNST was more strongly coupled with the salience network during approach (but no differential functional connectivity was detected when comparing approach vs. retreat). Finally, the PAG is another important brain region involved in threat processing (Bandler and Shipley, 1994). Here, we detected increased functional correlations between the right/left PAG with the salience network during approach; we only detected the left PAG as more strongly connected during approach vs. retreat. Note also that the interactions between several subcortical structures (left BNST, right PAG, and right habenula), and the salience network exhibited temporal properties and decreasing cohesion was observed as the circles moved away from each other during retreat.

In conclusion, in the present paper, we employed intersubject network analysis, which allows the investigation of network-level properties “across brains.” We found that threat altered network cohesion across the salience, executive, and task-negative networks, as well as subcortical regions. For example, cohesion increased within the salience network during approach relative to retreat. Functional connections between several subcortical regions and the salience network also increased during approach vs. retreat. The regions included the PAG, habenula, and amygdala, showing that the latter region is involved under conditions of relatively prolonged and uncertain threat, and not only linked to phasic stimuli. The BNST was functionally linked to regions of the salience network during approach, but we did not detect differential engagement when compared to retreat. Furthermore, cohesion within and between networks changed dynamically as threat imminence increased or decreased. In particular, salience-network cohesion increased during approach and decreased during retreat. Taken together, our findings unraveled dynamic properties of large-scale networks while threat levels varied continuously. The results demonstrate the potential of characterizing emotional processing at the level of distributed networks, and not simply at the level of evoked responses in specific brain regions. In particular, periods during which anxious anticipation waxes and wanes are paralleled by changes to brain network organization.

## Acknowledgements

The authors acknowledge funding from the National Institute of Mental Health (MH071589). We thank Srikanth Padmala and Elizabeth Redcay for feedback on the manuscript. We would like to thank Brenton McMenamin for paradigm development, Christian Meyer and Dan Levitas for data collection, Jason Smith for help with processing scripts, and Nicole Friedman and Jessica Berman for help with subject recruitment. The authors also acknowledge the Behavioral and Social Sciences College, University of Maryland, high performance computing resources (http://bsos.umd.edu/oacs/bsos-high-performance) made available for conducting the research reported in this paper.

## Author contributions

M.N. and L.P designed research; M.N analyzed the data together with L.P.; J.K. provided algorithms; L.P and M.N. wrote the paper.

## References

Alvarez, RP, Chen G, Bodurka J, Kaplan R, Grillon C (2011) Phasic and sustained fear in humans elicits distinct patterns of brain activity. NeuroImage 55:389–400.

Ames DL, Honey CJ, Chow, MA, Todorov A, Hasson U (2015) Contextual alignment of cognitive and neural dynamics. Journal of cognitive neuroscience.

Avants BB, Tustison NJ, Song G, Cook Pa, Klein A, Gee JC (2011) A reproducible evaluation of ANTs similarity metric performance in brain image registration. NeuroImage 54:2033–2044.

Avery SN, Clauss JA, Winder DG, Woodward N, Heckers S, Blackford JU (2014) BNST neurocircuitry in humans. Neuroimage 91:311–323.

Bandler R, Shipley MT (1994) Columnar organization in the midbrain periaqueductal gray: modules for emotional expression? Trends Neurosci 17:379–389.

Bannerman D, Rawlins J, McHugh S, Deacon R, Yee B, Bast T, Zhang W-N, Pothuizen H, Feldon J (2004) Regional dissociations within the hippocampus—memory and anxiety. Neuroscience & Biobehavioral Reviews 28:273–283.

Bullmore E, Sporns O (2009) Complex brain networks: graph theoretical analysis of structural and functional systems. Nature Reviews Neuroscience 10:186–198.

Choi JM, Padmala S, Pessoa L (2012) Impact of state anxiety on the interaction between threat monitoring and cognition. NeuroImage 59:1912–1923.

Cohen MS (1997) Parametric analysis of fMRI data using linear systems methods. NeuroImage 6:93–103.

Cox RW (1996) AFNI: Software for analysis and visualization of functional magnetic resonance neuroimages. Computers and Biomedical Research 29:162–173.

Davis M, Shi C (1999) The extended amygdala: are the central nucleus of the amygdala and the bed nucleus of the stria terminalis differentially involved in fear versus anxiety? Ann N Y Acad Sci 877:281–291.

Davis M, Walker DL, Miles L, Grillon C (2010a) Phasic vs sustained fear in rats and humans: role of the extended amygdala in fear vs anxiety. Neuropsychopharmacology 35:105–135.

Davis M, Walker DL, Miles L, Grillon C (2010b) Phasic vs sustained fear in rats and humans: role of the extended amygdala in fear vs anxiety. Neuropsychopharmacology 35:105–135.

Diedrichsen J, Balsters JH, Flavell J, Cussans E, Ramnani N (2009) A probabilistic MR atlas of the human cerebellum. Neuroimage 46:39–46.

Feinberg DA, Moeller S, Smith SM, Auerbach E, Ramanna S, Glasser MF, Miller KL, Ugurbil K, Yacoub E (2010) Multiplexed echo planar imaging for sub-second whole brain FMRI and fast diffusion imaging. PloS one 5:e15710.

Fox AS, Kalin NH (2014) A translational neuroscience approach to understanding the development of social anxiety disorder and its pathophysiology. American Journal of Psychiatry 171:1162–1173.

Fox AS, Oler JA, Tromp DP, Fudge JL, Kalin NH (2015) Extending the amygdala in theories of threat processing. Trends in neurosciences 38:319–329.

Fox E, Russo R, Georgiou GA (2005) Anxiety modulates the degree of attentive resources required to process emotional faces. Cognitive, Affective, & Behavioral Neuroscience 5:396–404.

Greve DN, Fischl B (2009) Accurate and robust brain image alignment using boundary-based registration. Neuroimage 48:63–72.

Grupe DW, Nitschke JB (2013) Uncertainty and anticipation in anxiety: an integrated neurobiological and psychological perspective. Nat Rev Neurosci 14:488–501.

Grupe DW, Oathes DJ, Nitschke JB (2013) Dissecting the anticipation of aversion reveals dissociable neural networks. Cereb Cortex 23:1874–1883.

Hasson U, Nir Y, Levy I, Fuhrmann G, Malach R (2004) Intersubject synchronization of cortical activity during natural vision. Science (New York, NY) 303:1634–1640.

Hermans EJ, van Marle HJ, Ossewaarde L, Henckens MJ, Qin S, van Kesteren MT, Schoots VC, Cousijn H, Rijpkema M, Oostenveld R, Fernandez G (2011) Stress-related noradrenergic activity prompts large-scale neural network reconfiguration. Science 334:1151–1153.

Hikosaka O (2010) The habenula: from stress evasion to value-based decision-making. Nature Reviews Neuroscience 11:503–513.

Iglesias JE, Liu C-Y, Thompson PM, Tu Z (2011) Robust brain extraction across datasets and comparison with publicly available methods. IEEE transactions on medical imaging 30:1617–1634.

Krauth A, Blanc R, Poveda A, Jeanmonod D, Morel A, Székely G (2010) A mean three-dimensional atlas of the human thalamus: generation from multiple histological data. Neuroimage 49:2053–2062.

Lahnakoski JM, Glerean E, Jääskeläinen IP, Hyönä J, Hari R, Sams M, Nummenmaa L (2014) Synchronous brain activity across individuals underlies shared psychological perspectives. Neuroimage 100:316–324.

McMenamin BW, Langeslag SJ, Sirbu M, Padmala S, Pessoa L (2014) Network Organization Unfolds over Time during Periods of Anxious Anticipation. The Journal of Neuroscience 34:11261–11273.

Menon V, Uddin LQ (2010) Saliency, switching, attention and control: a network model of insula function. Brain Struct Funct 214:655–667.

Mobbs D, Yu R, Rowe JB, Eich H, FeldmanHall O, Dalgleish T (2010) Neural activity associated with monitoring the oscillating threat value of a tarantula. Proceedings of the National Academy of Sciences of the United States of America 107:20582–20586.

Nacewicz B.M. A, A.L., Kalin, N.H. & Davidson R.J. (2014) The neurochemical underpinnings of human amygdala volume including subregional contributions. In: Society of Biological Psychiatry New York.

Najafi M, Pessoa L (2016) Studying intersubject networks and standard graph measures during dynamic threat processing. In: Program No 36904/NNN442016 Neuroscience Meeting Planner, vol.Online San Diego, CA: Society for Neuroscience, 2016.

Neter J, Kutner MH, Nachtsheim CJ, Wasserman W (1996) Applied linear statistical models (4th edition). Chicago: Irwin.

Newman M (2010a) Networks: An Introduction. New York City: Oxford University Press.

Newman MEJ (2010b) Networks: An Introduction.

Nummenmaa L, Glerean E, Viinikainen M, Jääskeläinen IP, Hari R, Sams M (2012) Emotions promote social interaction by synchronizing brain activity across individuals. Proceedings of the National Academy of Sciences 109:9599–9604.

Nummenmaa L, Saarimäki H, Glerean E, Gotsopoulos A, Jääskeläinen IP, Hari R, Sams M (2014)Emotional speech synchronizes brains across listeners and engages large-scale dynamic brain networks. Neuroimage 102:498–509.

Pessoa L (in press) A Network Model of the Emotional Brain. Trends in Cognitive Sciences.

Pessoa L, McMenamin B (2016) Dynamic Networks in the Emotional Brain. The Neuroscientist 1073858416671936.

Poldrack RA, Baker CI, Durnez J, Gorgolewski KJ, Matthews PM, Munafò MR, Nichols TE, Poline J-B, Vul E, Yarkoni T (2017) Scanning the horizon: towards transparent and reproducible neuroimaging research. Nature Reviews Neuroscience.

Power JD, Schlaggar BL, Lessov-Schlaggar CN, Petersen SE (2013) Evidence for hubs in human functional brain networks. Neuron 79:798–813.

Pruessner JC, Dedovic K, Khalili-Mahani N, Engert V, Pruessner M, Buss C, Renwick R, Dagher A, Meaney MJ, Lupien S (2008) Deactivation of the limbic system during acute psychosocial stress: evidence from positron emission tomography and functional magnetic resonance imaging studies. Biol Psychiatry 63:234–240.

Riedel MC, Ray KL, Dick AS, Sutherland MT, Hernandez Z, Fox PM, Eickhoff SB, Fox PT, Laird AR (2015) Meta-analytic connectivity and behavioral parcellation of the human cerebellum. Neuroimage 117:327–342.

Roy M, Shohamy D, Daw N, Jepma M, Wimmer GE, Wager TD (2014) Representation of aversive prediction errors in the human periaqueductal gray. Nature neuroscience 17:1607–1612.

Seeley WW, Menon V, Schatzberg AF, Keller J, Glover GH, Kenna H, Reiss AL, Greicius MD (2007) Dissociable intrinsic connectivity networks for salience processing and executive control. J Neurosci 27:2349–2356.

Shackman AJ, Fox AS (2016) Contributions of the Central Extended Amygdala to Fear and AnxietyContributions of the Central Extended Amygdala to Fear and Anxiety. Journal of neuroscience 36:8050–8063.

Shattuck DW, Leahy RM (2002) BrainSuite: an automated cortical surface identification tool. Medical image analysis 6:129–142.

Simony E, Honey CJ, Chen J, Lositsky O, Yeshurun Y, Wiesel A, Hasson U (2016) Dynamic reconfiguration of the default mode network during narrative comprehension. Nature communications 7

Somerville LH, Whalen PJ, Kelley WM (2010) Human bed nucleus of the stria terminalis indexes hypervigilant threat monitoring. Biological Psychiatry 68:416–424.

Stephens GJ, Silbert LJ, Hasson U (2010) Speaker–listener neural coupling underlies successful communication. Proceedings of the National Academy of Sciences 107:14425–14430.

Theiss JD, Ridgewell C, McHugo M, Heckers S, Blackford JU (2017) Manual segmentation of the human bed nucleus of the stria terminalis using 3T MRI. Neuroimage 146:288–292.

Torrisi S, O’connell K, Davis A, Reynolds R, Balderston N, Fudge JL, Grillon C, Ernst M (2015) Resting state connectivity of the bed nucleus of the stria terminalis at ultra-high field. Human brain mapping 36:4076–4088.

Tovote P, Fadok JP, Lüthi A (2015) Neuronal circuits for fear and anxiety. Nature Reviews Neuroscience 16:317–331.

Van Dijk KR, Sabuncu MR, Buckner RL (2012) The influence of head motion on intrinsic functional connectivity MRI. Neuroimage 59:431–438.

Wager TD, Waugh CE, Lindquist M, Noll DC, Fredrickson BL, Taylor SF (2009) Brain mediators of cardiovascular responses to social threat. Neuroimage 47:821–835.

Yeo BT, Krienen FM, Sepulcre J, Sabuncu MR, Lashkari D, Hollinshead M, Roffman JL, Smoller JW, Zollei L, Polimeni JR, Fischl B, Liu H, Buckner RL (2011) The organization of the human cerebral cortex estimated by intrinsic functional connectivity. Journal of Neurophysiology 106:1125–1165.

